# NMNAT1 Binding at Promoters and Enhancers Couples NAD^+^ Synthesis to RNA Polymerase II Engagement

**DOI:** 10.1101/2025.04.30.651499

**Authors:** Yessenia Cedeño-Cedeño, Samuel J. Taylor, Matthew J. Gamble, Ulrich Steidl, Robert A. Coleman

## Abstract

Gene expression relies on transcriptional bursts driven by dynamic chromatin-modifying enzymes that depend on metabolites like NAD⁺. Depletion of NAD^+^ contributes to cancer, metabolic disorders, and aging, emphasizing the significance of tight regulation of NAD^+^ production in the cell. NMNAT1, a nuclear NAD⁺-synthetase enzyme, supports chromatin-modifying enzymes such as PARP1 and SIRT1; however, its direct role in transcriptional regulation remains unclear. Using integrated multi-omics, we present the first high-resolution, genome-wide study of NMNAT1’s regulatory functions. We demonstrate that NMNAT1 binds to the promoters and enhancers of actively transcribed genes involved in DNA replication, cell cycle progression, and chromatin regulation. RNA-seq and CUT&Tag analyses indicate reduced RNA Polymerase II occupancy at downregulated genes in NMNAT1 knockout cells, implicating NMNAT1 in transcriptional activation through Pol II engagement. These findings position NMNAT1 as a key node linking localized NAD⁺ production to gene-specific transcription, offering new insights into metabolic regulation of gene expression.

## INTRODUCTION

Gene expression is an energetically demanding and highly dynamic process driven by transcriptional bursts and rapid chromatin remodeling^1^. These dynamics are partially regulated by chromatin-modifying enzymes, which change chromatin accessibility and modify histones or the transcriptional machinery to control gene expression^2–5^. Many metabolite-dependent enzymes require cofactors such as nicotinamide adenine dinucleotide (NAD⁺) to support their enzymatic functions^6–10^. Studies showing the presence of nuclear metabolic enzymes at specific locations on the genome highlight that local production of nuclear metabolites may directly impact chromatin states and transcriptional activity^11–13^.

NAD⁺ is a central coenzyme with essential roles in redox metabolism, DNA repair, and gene regulation^14–20^. Cellular NAD⁺ levels decline with age and are dysregulated in multiple diseases, including cancer and metabolic disorders, underscoring its importance in maintaining genomic and transcriptional homeostasis^6,17,19,21–26^. Although NAD⁺ can diffuse into the nucleus from other compartments, accumulating evidence suggests that nuclear-localized NAD⁺ synthesis is spatially coordinated with gene regulatory processes^20^.

The enzyme nicotinamide mononucleotide adenylyltransferase 1 (NMNAT1) catalyzes the final step of the nuclear NAD⁺ salvage pathway^27^. While NMNAT1 is traditionally considered a metabolic enzyme, emerging studies suggest a regulatory role in chromatin biology^7,28–32^. Low-resolution ChIP-chip and biochemical analyses have shown that NMNAT1 colocalizes with NAD⁺-dependent chromatin-modifying enzymes such as PARP1 and SIRT1^28,29,31–33^. In addition, NMNAT1’s depletion is known to disrupt PARP1 and SIRT1 enzymatic activity and gene expression of target genes. In vitro studies further indicate that NMNAT1 can modulate PARP1 activity via allosteric interactions, independent of its catalytic activity^34^. However, the direct genomic binding profile of NMNAT1 and its broader role in transcriptional regulation remain uncharacterized.

Here, we provide the first high-resolution, genome-wide analysis of NMNAT1 occupancy and function using integrative multi-omics approaches. CUT&Tag profiling revealed that NMNAT1 is enriched at promoter and enhancer elements of actively transcribed genes involved in DNA replication, cell cycle progression, and chromatin regulation. Integration of transcriptomic and genomic localization assays revealed that NMNAT1 knockout leads to the downregulation of direct target genes and a corresponding decrease in RNA Polymerase II occupancy at their core promoters. These findings suggest that NMNAT1 promotes transcriptional activation by potentially facilitating RNA-Pol II engagement at select genes.

Our findings establish NMNAT1 as a crucial regulatory point linking local nuclear NAD⁺ synthesis to transcriptional output. This research expands our understanding of the roles played by nuclear metabolic enzymes that selectively bind the genome. It offers new insights into how local metabolite production interacts with the fundamental transcriptional machinery to regulate gene expression programs, potentially affecting disease-related pathways such as cancer, metabolic disorders, and age-related diseases.

## RESULTS

### NMNAT1 Knockout Disrupts NAD⁺ Homeostasis and Impairs NAD⁺-Dependent Chromatin Modifier Activity

NMNAT1 catalyzes the final step in the nuclear NAD⁺ salvage pathway and is thought to locally regulate the availability of NAD⁺ for nuclear enzymes^35–37^. While NAD⁺ depletion has been linked to genomic instability, metabolic dysfunction, and age-related decline, the specific contribution of NMNAT1 to nuclear metabolism and chromatin regulation remains poorly defined^6,15,17,21,24,38,39^. To investigate NMNAT1 function, we knocked out NMNAT1 (NMNAT1^KO^) in human U-2OS cells via CRISPR/Cas9 targeting of exons 3–5 (Figure 1A)^33^. Western blot analysis confirmed loss of NMNAT1 protein expression in NMNAT1^KO^ cells (Figure 1B).

**Figure 1.**
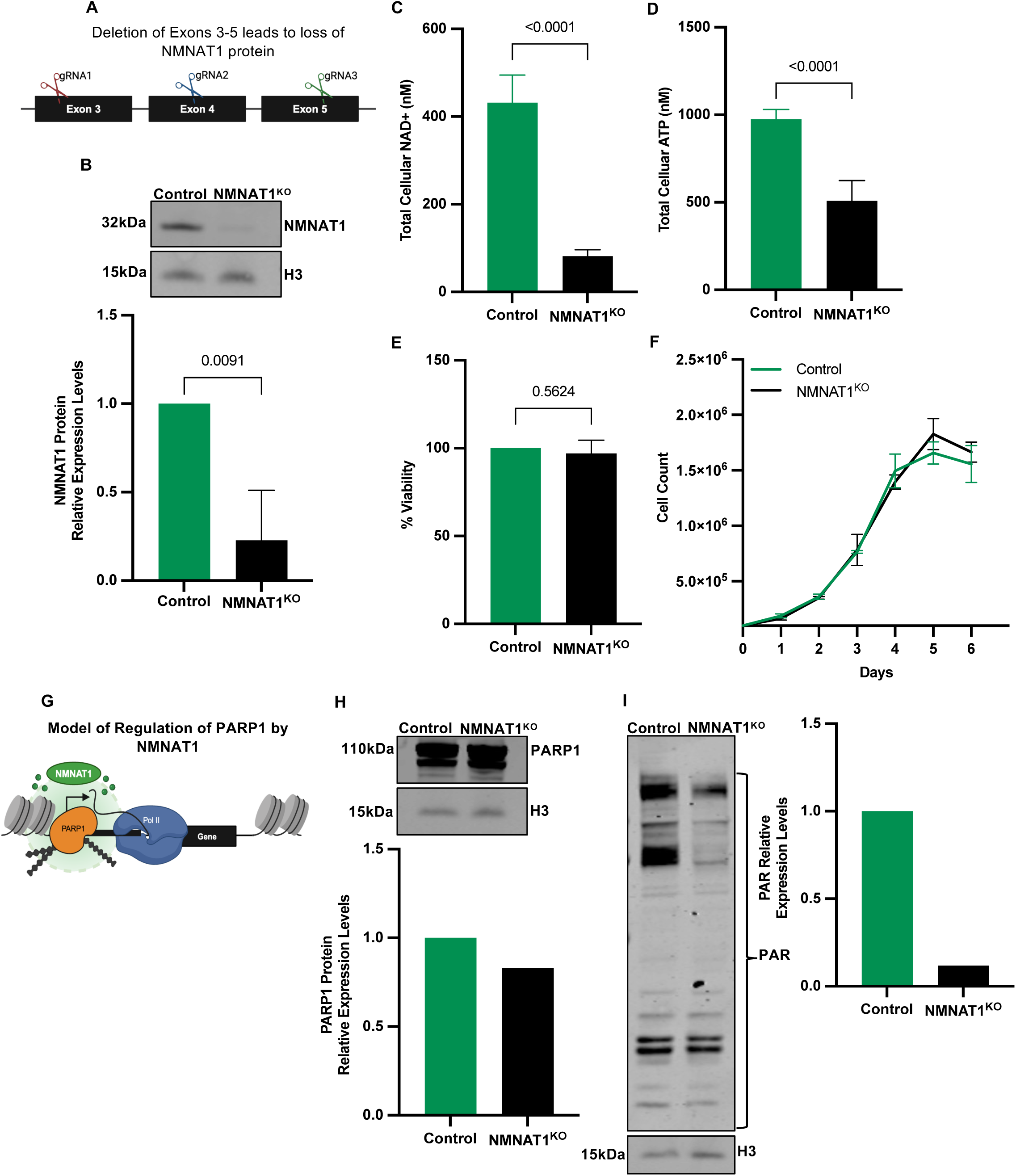
NMNAT1 Knockout Disrupts NAD⁺ Homeostasis and Impairs NAD⁺-Dependent Chromatin Modifier Activity. (A) Schematic of CRISPR/Cas9 strategy targeting exons 3–5 of NMNAT1 in U-2OS cells to generate a frameshift mutation and truncated protein. (B) Western blot confirming NMNAT1 depletion in NMNAT1^KO^ cells. Quantification normalized to H3 is shown (mean ± SEM, n = 3; p < 0.01, unpaired t-test). (C) Total NAD⁺ levels are significantly reduced in NMNAT1^KO^ cells as measured by NAD⁺/NADH assay (mean ± SEM, n = 3; p < 0.01, unpaired t-test). (D) ATP levels are significantly decreased in NMNAT1^KO^ cells (mean ± SEM, n = 5; p < 0.01, unpaired t-test). (E) Calcein-AM viability assay reveals no significant change in NMNAT1^KO^ cell viability (mean ± SEM, n = 3). (F) Growth curve analysis shows comparable proliferation rates between NMNAT1^KO^ and control cells over five days (mean ± SEM, n = 2). (G) Model of NMNAT1-regulated transcription via NAD⁺-dependent PARP1 activation and promoter-associated PARylation. Adapted from Zhang et al., *J. Biol. Chem.* (2009, 2012). (H) PARP1 protein levels remain unchanged in NMNAT1^KO^ cells. Quantification shown (mean ± SEM, n = 1; p < 0.01, unpaired t-test). (I) Global PAR levels are significantly reduced in NMNAT1^KO^ cells, indicating loss of PARP1 activity (mean ± SEM, n = 1; p < 0.01, unpaired t-test).

A colorimetric NAD⁺/NADH assay indicated an approximately 75% decrease in total cellular NAD⁺ levels in NMNAT1^KO^ cells compared to controls (Figure 1C). This finding aligns with previous studies on NMNAT1 silencing in other contexts^28,33^. We measured total ATP levels since NAD^+^ metabolism is intertwined with ATP production. We found that ATP levels are significantly decreased in NMNAT1^KO^ cells (Figure 1D). Despite the substantial NAD⁺ and ATP depletion, NMNAT1^KO^ cells remained viable as determined by Calcein-AM fluorescence (Figure 1E) with normal growth rates (Figure 1F).

Considering that chromatin-modifying enzymes depend on NAD⁺, we assessed how the loss of NMNAT1 affects poly-(ADP-ribose) polymerase 1 (PARP1), a crucial NAD⁺-dependent regulator of chromatin (Figure 1G). While PARP1 protein levels remained constant in NMNAT1^KO^ cells (Figure 1H), the overall levels of poly-(ADP-ribose) (PAR), an indicator of PARP1’s enzymatic activity, were significantly lower (Figure 1I). These results indicate that NAD^+^ derived from NMNAT1 is essential for maintaining PARP1’s functional activity. This data establishes NMNAT1 as a central metabolic regulator of total cellular NAD⁺ and NAD⁺-dependent chromatin-modifying enzyme activity. The loss of NMNAT1 leads to widespread NAD⁺ depletion and impaired PARP1 activity, supporting a critical role for NMNAT1 in linking nuclear metabolism to chromatin regulation and transcriptional control.

### Loss of NMNAT1 Reprograms Transcriptional Networks Governing Nuclear and Extracellular Functions

Given NMNAT1’s essential role in maintaining nuclear NAD⁺ levels and supporting chromatin-modifying enzyme activity, we next investigated how its loss alters global gene expression. We performed bulk RNA sequencing in U-2OS NMNAT1^KO^ and control cells (Figure 2). Principal component analysis (PCA) revealed distinct clustering of NMNAT1^KO^ and control transcriptomes, with PC1 and PC2 accounting for 87% and 8% of the total variance, respectively (Figure 2A), indicating robust transcriptional reprogramming following NMNAT1 depletion.

**Figure 2.**
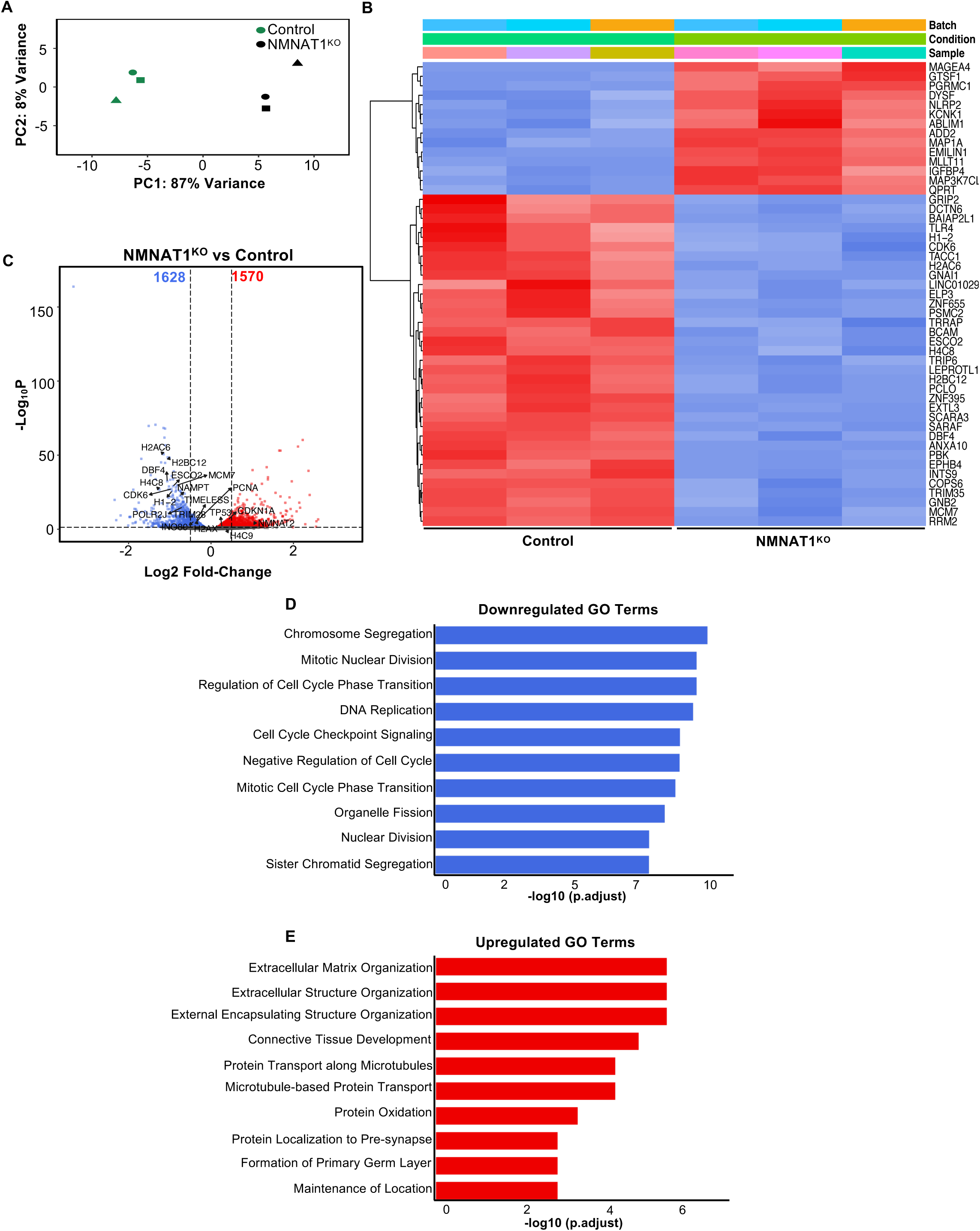
NMNAT1 Loss Reprograms Transcriptional Networks Governing Nuclear and Extracellular Functions. (A) Principal component analysis (PCA) of VST-transformed RNA-seq data shows distinct clustering of NMNAT1KO and control samples (PC1: 87%, PC2: 8%). (B) Heatmap of DEGs illustrating hierarchical clustering of transcriptomes between NMNAT1KO and control cells. Expression values are scaled row-wise. (C) Volcano plot showing log₂ fold change versus −log₁₀ adjusted *p*-value. Significant DEGs (adjusted *p* < 0.05) are colored red (up) and blue (down). (D) GO analysis of downregulated DEGs reveals enrichment for DNA replication, chromosome segregation, and cell cycle pathways. GO terms are plotted based on −log10 adjusted p-values. (E) GO analysis of upregulated DEGs identifies enrichment in extracellular matrix organization, protein transport, and vesicle-mediated trafficking. GO terms are plotted based on −log10 adjusted p-values.

Hierarchical clustering of normalized gene expression profiles further highlighted distinct transcriptomic states between conditions (Figure 2B). Differential expression analysis identified 3,185 significantly altered genes (adjusted *p* < 0.05), with 1,620 genes upregulated and 1,565 downregulated in NMNAT1^KO^ cells (Figure 2C, Table S1). These findings suggest that NMNAT1 is broadly required for transcriptional homeostasis in U-2OS cells.

To identify affected biological processes, we performed Gene Ontology (GO) enrichment analysis on the differentially expressed gene sets. Downregulated genes were significantly enriched in pathways related to DNA replication, cell cycle progression, and chromatin organization, processes closely associated with NAD⁺-dependent enzymes such as PARP1 and SIRT1 (Figure 2D). Conversely, upregulated genes were enriched in pathways related to extracellular matrix organization, protein transport, and vesicle-mediated trafficking (Figure 2E), which are processes associated with other members of the PARP family (e.g., PARP2)^40^.

Among the downregulated genes, we observed a coordinated decrease in genes involved in DNA replication (e.g., *ORC5, MCM7, ESCO2*), transcription (e.g., *POLR2J, TAF6, TRIM28*), and chromatin structure (e.g., *H1-2*, *H2AC6, H2BC12*, *H4-8*). These genes represent key components of essential nuclear processes, reinforcing that NMNAT1 loss disrupts core transcriptional programs. Intriguingly, the expression of NAMPT, the metabolic enzyme that provides a key metabolite for all NMNATs in the cell, was downregulated upon knockout of NMNAT1^41^. This result suggests a key feedforward loop whereby NMNAT1 regulates not only nuclear but also global NAD^+^ levels as well. Together, these data demonstrate that the knockout of NMNAT1 and the resulting depletion of nuclear NAD^+^ lead to gene-specific transcriptional dysregulation, particularly affecting genes that are essential for genome maintenance and cell proliferation. These findings support a model in which nuclear metabolic enzymes, such as NMNAT1, regulate gene expression by sustaining the activity of NAD⁺-dependent chromatin modifiers and preserving nuclear transcriptional programs.

### NMNAT1 Selectively Localizes to Promoter and Enhancer Regions Through Transcription Factor–Linked Recruitment

The specific transcriptional changes observed in NMNAT1 knockout cells suggest that NMNAT1 may exert direct regulatory effects at the chromatin level. While prior low-resolution ChIP-chip studies reported NMNAT1 enrichment at promoters of PARP1 and SIRT1 target genes (e.g., *ATXN10, NAT1, SOCS2, PEG10, TMSNB, and NELL2*), these studies lacked genome-wide resolution^31,32^. As a result, the genomic binding landscape of NMNAT1 remains only partially defined. To better define NMNAT1 occupancy, we conducted CUT&Tag analysis of NMNAT1 localization in U-2OS cells (Figure 3A).

**Figure 3.**
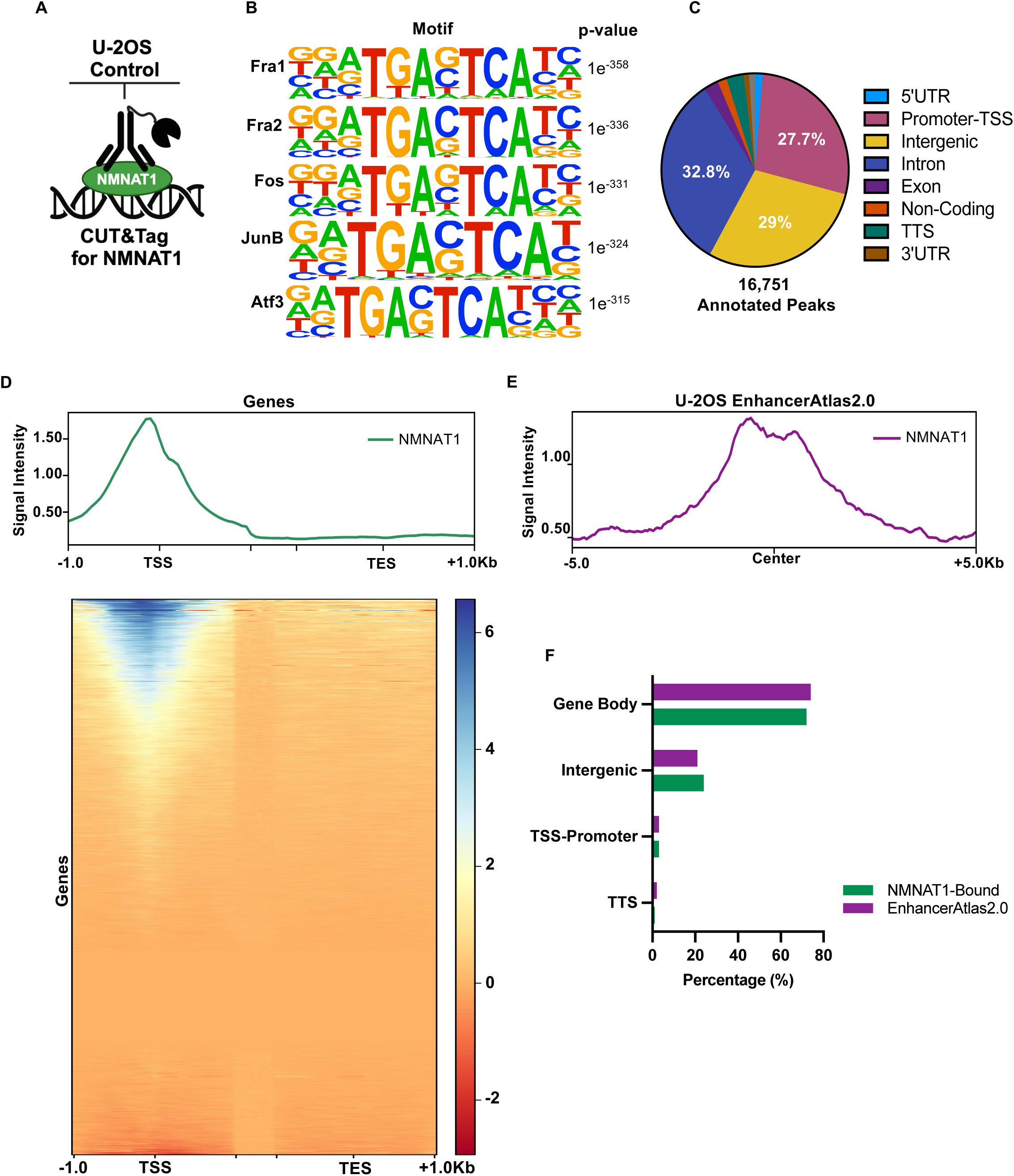
NMNAT1 Selectively Localizes to Promoter and Enhancer Regions via Transcription Factor–Mediated Recruitment. (A) Schematic of the CUT&Tag workflow to profile NMNAT1 chromatin binding in U-2OS cells. The signal from NMNAT1^KO^ cells was subtracted to define high-confidence peaks. (B) Motif analysis of NMNAT1-bound regions performed using HOMER shows enrichment of AP-1 transcription factor motifs, including Fra1/2, Fos, JunB, and Atf3. (C) Genomic distribution of 16,751 NMNAT1 peaks identified from CUT&Tag analysis. Peaks were annotated relative to gene features, including promoters (27.7%), introns (32.8%), and distal intergenic regions (29%). (D) Metaplot and heatmap of NMNAT1 signal centered on transcription start sites (TSS ±1 kb), showing non-uniform enrichment across promoter regions. (E) Metaplot of NMNAT1 signal centered on known U-2OS enhancers from EnhancerAtlas2.0. (F) Distribution (percentages) of NMNAT1-bound at known enhancers (U-2OS EnhancerAtlas2.0).

To identify high-confidence binding sites, we subtracted the background CUT&Tag signal obtained in NMNAT1 knockout cells from wild-type U-2OS CUT&Tag profiles (see Methods). We identified 16,752 NMNAT1 peaks genome-wide. To investigate possible recruitment mechanisms of NMNAT to target sites, we performed a de novo motif enrichment analysis using HOMER. We found that NMNAT1-bound regions were significantly enriched for AP-1 family transcription factor motifs, such as Fra1/2, Fos, JunB, and Atf3 (Figure 3B). ENCODE data reveal that these factors frequently co-occupy many of the same regions in other cell types. This supports a model where NMNAT1 is recruited to chromatin through select interactions with transcription factors.

NMNAT1 peaks were distributed across promoter regions (27.7%), introns (32.8%), and distal intergenic elements (29%) (Figure 3C). Notably, a substantial fraction of NMNAT1 peaks localized to promoter–TSS regions (±1 kb), implicating NMNAT1 in transcription initiation. Metaplots and heat maps centered on TSS regions showed that the NMNAT1 signal was strongly enriched near the TSS but was concentrated at a subset of genes, indicating selective promoter association and potential gene-specific regulatory activity (Figure 3D).

In addition to promoter localization, we discovered that NMNAT1 is enriched at enhancer elements (Figure 3E). Using enhancer annotations from EnhancerAtlas 2.0 specific to U-2OS cells, we identified a subset of 3,506 NMNAT1 peaks that overlapped with known cell-type– specific enhancers. These enhancer sites were distributed across intergenic (24%), promoter-TSS (3%), and gene body (73%) regions (Figure 3F). These findings suggest that NMNAT1 may contribute to enhancer-mediated transcriptional regulation, potentially facilitating long-range enhancer-promoter communication or modulating enhancer activity through local NAD^+^ synthesis. Together, these results present the first high-resolution, genome-wide chromatin occupancy map of NMNAT1 in human cells. They reveal that NMNAT1 selectively binds to promoter and enhancer regions, likely through transcription factor-mediated recruitment. They highlight its broader role in regulating proximal and distal gene regulatory elements through nuclear NAD⁺ metabolism.

### NMNAT1 Promotes Transcriptional Activation Through Selective Engagement at Promoter–TSS and Enhancer Regions

Having established that NMNAT1 localizes to promoter and enhancer elements throughout the genome, we next aimed to determine whether NMNAT1 binding directly influences transcriptional output. To address this, we performed an integrative analysis of NMNAT1 CUT&Tag and RNA-seq data in U-2OS cells (Figure 4A). We initially analyzed NMNAT1 enrichment among various classes of differentially expressed genes. Metaplot analysis indicates that NMNAT1 binds more prominently at the promoter–TSS regions of downregulated genes, in contrast to unchanged genes (Figure 4B). Conversely, NMNAT1 exhibited lower occupancy at genes upregulated by NMNAT1^KO^, indicating that NMNAT1 functions to help positively regulate transcription at select genes.

**Figure 4.**
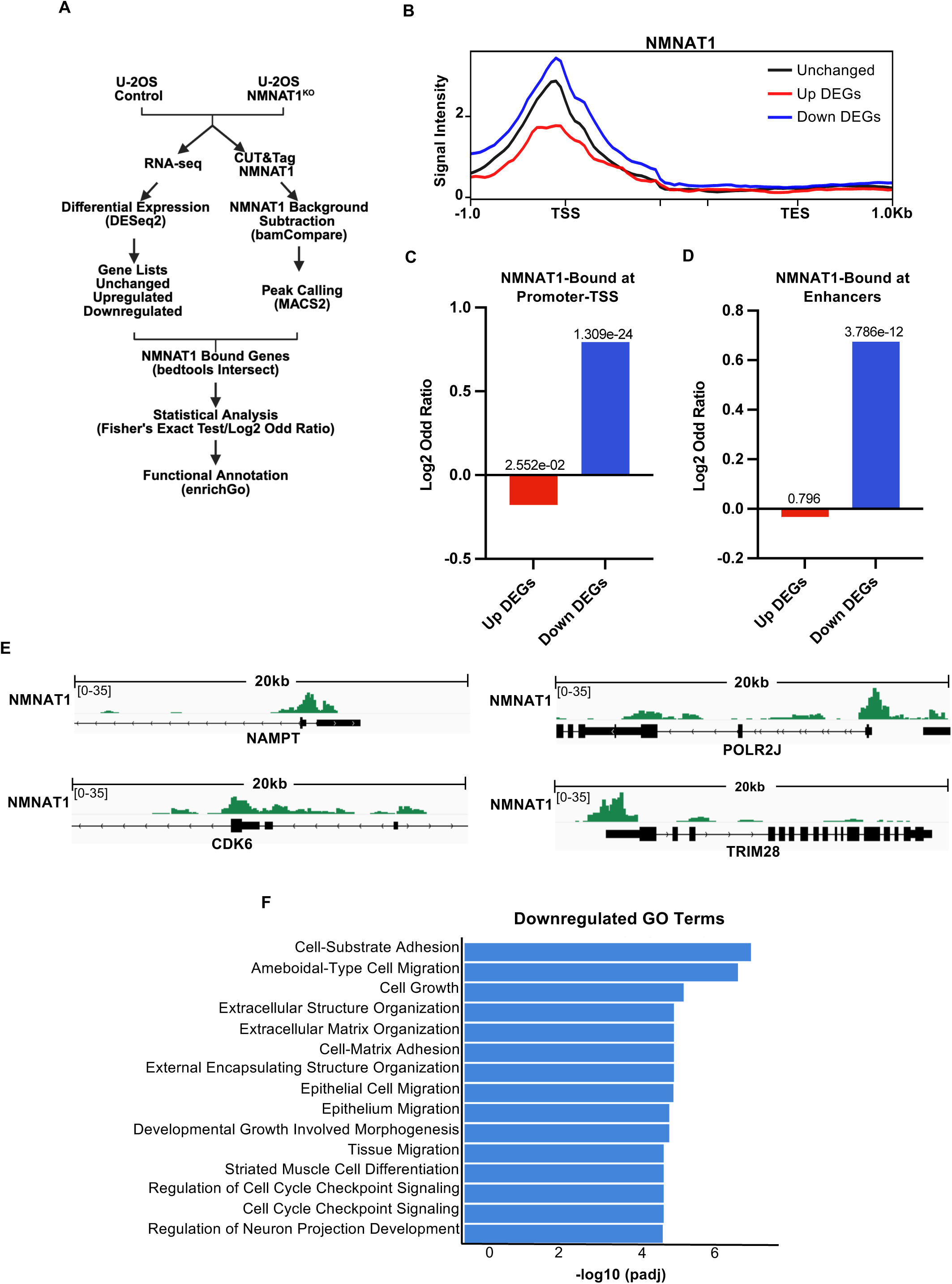
NMNAT1 Occupancy at Promoter–TSS and Enhancer Regions Correlates with Transcriptional Activation of Downregulated Genes. (A) Schematic of the integrative analysis pipeline combining NMNAT1 CUT&Tag with RNA-seq data to assess the relationship between chromatin occupancy and gene expression in U-2OS cells. (B) Metaplot of NMNAT1 signal intensity at ±1 kb of the transcription start site (TSS) and transcription end site (TES) across unchanged (black), upregulated (red), and downregulated (blue) genes. NMNAT1 shows the highest enrichment at the TSS of downregulated genes. (C) Log₂ odds ratio plot quantifying the enrichment of NMNAT1 binding at the promoter-TSS of downregulated genes (blue) compared to unchanged and upregulated genes (red). Significant enrichment of NMNAT1 peaks was observed at downregulated genes (*p* = 1.309e^−24^) but not at upregulated genes (See also Figure S1 for complete statistical analysis). (D) Log₂ odds ratio plot quantifying the enrichment of NMNAT1 binding at enhancers of downregulated genes (blue) compared to unchanged and upregulated genes (red). Significant enrichment of NMNAT1-enhancer bound was observed at downregulated genes (*p* = 3.786e^−12^) but not at upregulated genes (See also Figure S1 for complete statistical analysis). (E) Genome browser tracks of NMNAT1 CUT&Tag signal in U-2OS control subtracted from CUT&Tag signal in NMNAT1^KO^ cells at selected target gene promoters. (F) Gene Ontology (GO) analysis of NMNAT1-bound downregulated genes. Significantly enriched terms include cell cycle regulation, DNA replication, and extracellular matrix organization. GO terms are ranked by −log₁₀ adjusted *p*-value.

To determine whether NMNAT1 binding on differentially regulated genes is statistically significant, we performed a Fisher’s Exact Test and calculated log₂ odds ratios comparing NMNAT1 peak overlap among differentially expressed gene sets. This analysis revealed a significant enrichment of NMNAT1 peaks at the promoter-TSS of downregulated genes and depletion of NMNAT1 on upregulated genes (Figure 4C and S1). Furthermore, NMNAT1 is also significantly enriched at known enhancers of downregulated genes (Figure 4D). Visualization of CUT&Tag data on individual genes confirmed the presence of NMNAT1 at the promoters and enhancers of downregulated target genes (Figure 4E). GO enrichment analysis of NMNAT1-bound downregulated genes revealed functional categories essential for cellular homeostasis, including cell cycle regulation, DNA replication, cell growth, and extracellular matrix organization (Figure 4F). These findings indicate that NMNAT1 actively enhances gene expression by targeting the promoter and enhancer regions of crucial regulatory genes. Instead of affecting the entire transcriptome, NMNAT1 seems to specifically interact with a significant subset of genes, linking localized NAD⁺ production to transcriptional activation, thereby supporting vital cellular pathways.

### NMNAT1 Is Required for RNA Polymerase II Engagement at Promoter–TSS Regions

The selective transcriptional dysregulation observed in NMNAT1^KO^ cells, particularly at cell cycle and chromatin regulation genes, led us to investigate whether NMNAT1 influences engagement of the transcriptional machinery at our NAD⁺-regulated genes. Given its role in sustaining nuclear NAD⁺ and supporting PARP1 activity, we hypothesized that NMNAT1 depletion may disrupt promoter-proximal pause release of RNA-Pol II via a PARP1–NELF–dependent mechanism^30^ (Figure 5A). In such a scenario, we would expect RNA-Pol II signals to increase at the promoter regions of down-regulated genes.

**Figure 5.**
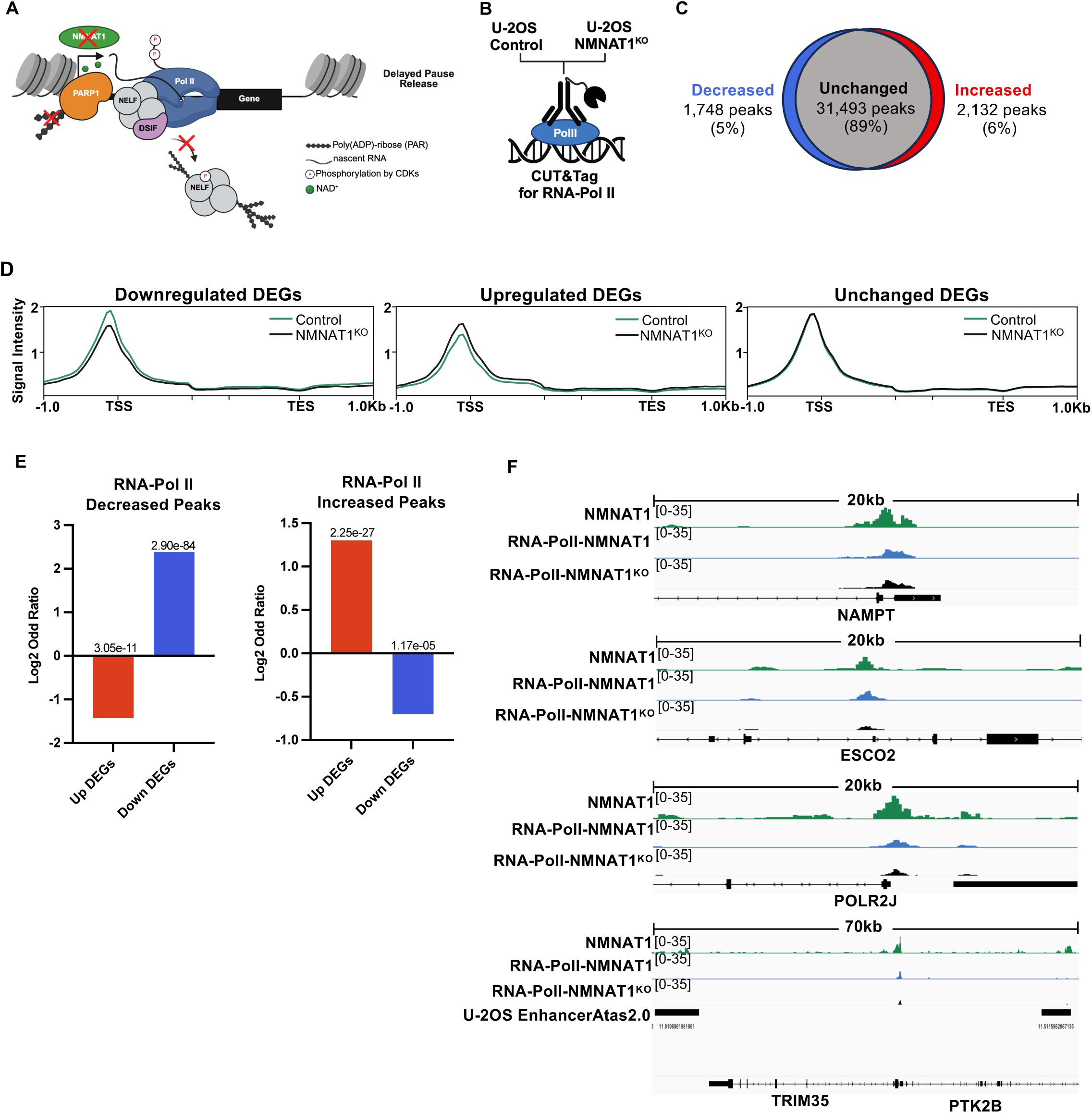
NMNAT1 Is Required for RNA Polymerase II Engagement at Promoters and Transcriptional Activation of Downregulated Genes. (A) Schematic model of NMNAT1 function in transcriptional regulation. NMNAT1 is proposed to support RNA-Pol II pausing and release via NAD⁺-dependent PARP1 activity and NELF, adding to the previous model^30^. (B) Schematic of CUT&Tag experimental workflow used to profile RNA-Pol II Ser5P occupancy in U-2OS control and NMNAT1^KO^ cells. (C) Differential peak analysis of 35,373 RNA-Pol II Ser5P peaks identifies 1,748 (5%) with decreased and 2,132 (6%) with increased occupancy in NMNAT1^KO^ cells relative to controls. Analysis performed using the GoodpeaksScript pipeline. (D) Metaplot of RNA-Pol II Ser5P signal across downregulated (left panel), upregulated (middle panel), and unchanged (right panel) DEGs. Reduced signal is observed specifically at downregulated genes in NMNAT1^KO^ cells. (E) Log₂ odds ratio plot quantifying the enrichment of RNA Polymerase II (Pol II) occupancy changes at downregulated (blue), unchanged, and upregulated (red) genes. Significant enrichment of RNA-Pol II-decreased peaks was observed at downregulated genes (p = 2.90e^−84^) but not at upregulated genes. Conversely, RNA-Pol II–increased peaks were significantly enriched at upregulated genes (p = 2.25e^−27^) but not at downregulated genes (see also Figure S2 for complete statistical analysis). (F) Genome browser tracks RNA-Pol II Ser5P occupancy at representative target genes in control and NMNAT1^KO^ U-2OS cells.

To test this hypothesis, we performed CUT&Tag profiling of RNA-Pol II phosphorylated at serine 5 of its C-terminal domain (Ser5P) in U-2OS control and NMNAT1^KO^ cells (Figure 5B). RNA-Pol II with Ser5P is enriched at TSS and reflects engaged, promoter-proximal RNA-Pol II. Differential peak analysis identified 35,373 RNA-Pol II Ser5P peaks genome-wide, of which 1,748 (5%) were decreased and 2,132 (6%) were increased upon NMNAT1 depletion (Figure 5C). We focused on differentially expressed genes (DEGs) identified from RNA-seq to examine the functional consequences of impaired RNA-Pol II occupancy. Metaplot analysis revealed a marked reduction in RNA-Pol II Ser5P signal at the promoters of downregulated genes in NMNAT1^KO^ cells (Figure 5D, left panel). Correspondingly, there was an increase in the RNA-Pol II Ser5P signal at the promoters of upregulated genes in NMNAT1^KO^ cells (Figure 5D, center panel). As a control, we also analyzed the RNA-Pol II Ser5P signal at unchanged genes and saw no difference in occupancy in NMNAT1^KO^ cells (Figure 5D, right panel). These findings suggest that NMNAT1 is required for productive Pol II engagement at specific transcriptionally active loci.

To quantify this relationship, we performed Fisher’s Exact Test and calculated log₂ odds ratios comparing RNA-Pol II Ser5P peak overlap across DEG categories. RNA-Pol II peaks were significantly enriched at downregulated DEGs relative to upregulated genes (Figure 5E and S2), further supporting a positive regulatory role for NMNAT1 in RNA-Pol II engagement at critical gene promoters. Visualization of CUT&Tag data on individual genes confirmed the reduction of RNA-Pol II at the promoters of downregulated target genes (Figure 5F). Together, these results reveal that NMNAT1 facilitates RNA Polymerase II recruitment or retention at promoter regions of key regulatory genes, likely through mechanisms that intersect with NAD⁺-dependent chromatin modification. While prior models have proposed a role for PARP1 in modulating transcriptional pausing via NELF, our findings suggest that NMNAT1 acts upstream of these pathways to enable proper transcriptional initiation and elongation.

## DISCUSSION

NMNAT1 is a member of an evolutionarily conserved family of enzymes responsible for NAD⁺ biosynthesis in bacteria, plants, and animals^36,42^. Despite its ancient origin and central role in cellular metabolism, the regulatory functions of NMNAT1 in gene expression have largely remained unexplored. Our research indicates that NMNAT1 acts as a metabolic catalyst while directly and selectively regulating the transcriptional output of select genes. We demonstrate that NMNAT1 knockout leads to a significant depletion of total cellular NAD⁺ and ATP levels. This considerable reduction in total cellular NAD⁺ is likely due to NMNAT1’s role in positively regulating the expression of NAMPT, which catalyzes the production of NMN, a metabolite utilized by all cellular NMNATs to produce NAD⁺. Therefore, NMNAT1 serves as a key nexus to regulate a cell’s response when there is an increased demand for NAD⁺ production during such processes as DNA repair.

Despite low levels of NAD⁺, U-2OS cells devoid of NMNAT1 maintain normal viability and proliferation under basal conditions, indicating metabolic resilience in this context^33^. In contrast, NMNAT1 depletion in the leukemia cell line MOLM13 impairs cell growth and viability, activating the p53 pathway and inducing cell cycle arrest^28^. These differences suggest that NMNAT1 function and NAD⁺ dependency may be cell-type or tissue-specific, potentially reflecting distinct metabolic demands or transcriptional programs. Nevertheless, even in metabolically resilient U-2OS cells, the loss of NMNAT1 alters transcriptional networks. Transcriptome profiling revealed specific gene expression changes, with significant downregulation of genes involved in DNA replication, cell cycle control, and chromatin organization. These findings underscore a key role for NMNAT1 in maintaining transcriptional homeostasis of core nuclear programs.

High-resolution CUT&Tag mapping has shown that NMNAT1 selectively localizes to the promoter and enhancer regions of actively transcribed genes. Motif analysis revealed that regions bound by NMNAT1 are rich in motifs from the AP-1 family of transcription factors, suggesting that NMNAT1 may be recruited to chromatin through interactions with sequence-specific transcription factors. AP-1 factors, including FOS, JUN, and ATF3, are recognized regulators of stress responses, proliferation, and differentiation programs^43–45^. This indicates that NMNAT1 could be associated with the transcriptional control of genes responsive to environmental or metabolic cues. Although prior research indicated NMNAT1 colocalizes with chromatin modifiers such as PARP1 and SIRT1^31,32^, this study is the first to detail its genome-wide occupancy and directly associate its binding with alterations in gene expression. Significantly, NMNAT1 enrichment was strongest at the transcription start sites and enhancers of downregulated genes, indicating its primary involvement in transcriptional activation instead of repression. These results define, for the first time, the genomic occupancy of NMNAT1 and establish a direct connection between its chromatin binding and transcriptional regulation.

Previous models have proposed that PARP1 facilitates the release of proximally paused RNA-Pol II through the PARylation of the NELF complex at specific genes^30^. Based on this model, we expected increased RNA-Pol II occupancy at our downregulated genes in cells lacking NMNAT1. However, we found that the depletion of NMNAT1 leads to a broad reduction in RNA-Pol II occupancy at transcription start sites of genes downregulated in NMNAT1^KO^ cells. This reduction indicates a defect in RNA Pol II recruitment or promoter-proximal engagement in the absence of NMNAT1. Our data suggest that PARP1, via NMNAT1’s local production of NAD⁺, likely targets additional proteins involved in regulating RNA-Pol II recruitment to genes. However, we cannot overlook the potential contribution of SIRT1, another NAD⁺-dependent enzyme and well-established regulator of transcription, which may also play a role in modulating RNA-Pol II engagement.

In support of this model, PARP1 has also been shown to PARylate transcription factors and components of the transcriptional machinery (e.g., pre-initiation factors including subunits of TFIID, TFIIB, TFIIE, TFIIF, TFIIH, and RNA-Pol I)^46–48^. In some cases, transcription factor activity, stability, and chromatin recruitment depend on PARP1 activity. However, the role of PARP1 in modulating the activity of many of these transcriptional regulators remains unknown. Our findings suggest that NMNAT1 may locally act upstream of RNA-Pol II pause release by facilitating the NAD⁺-dependent activation of PARP1, thereby coordinating chromatin remodeling, modulating transcription factors, and engagement of RNA-Pol II at select gene loci (Figure 6).

**Figure 6.**
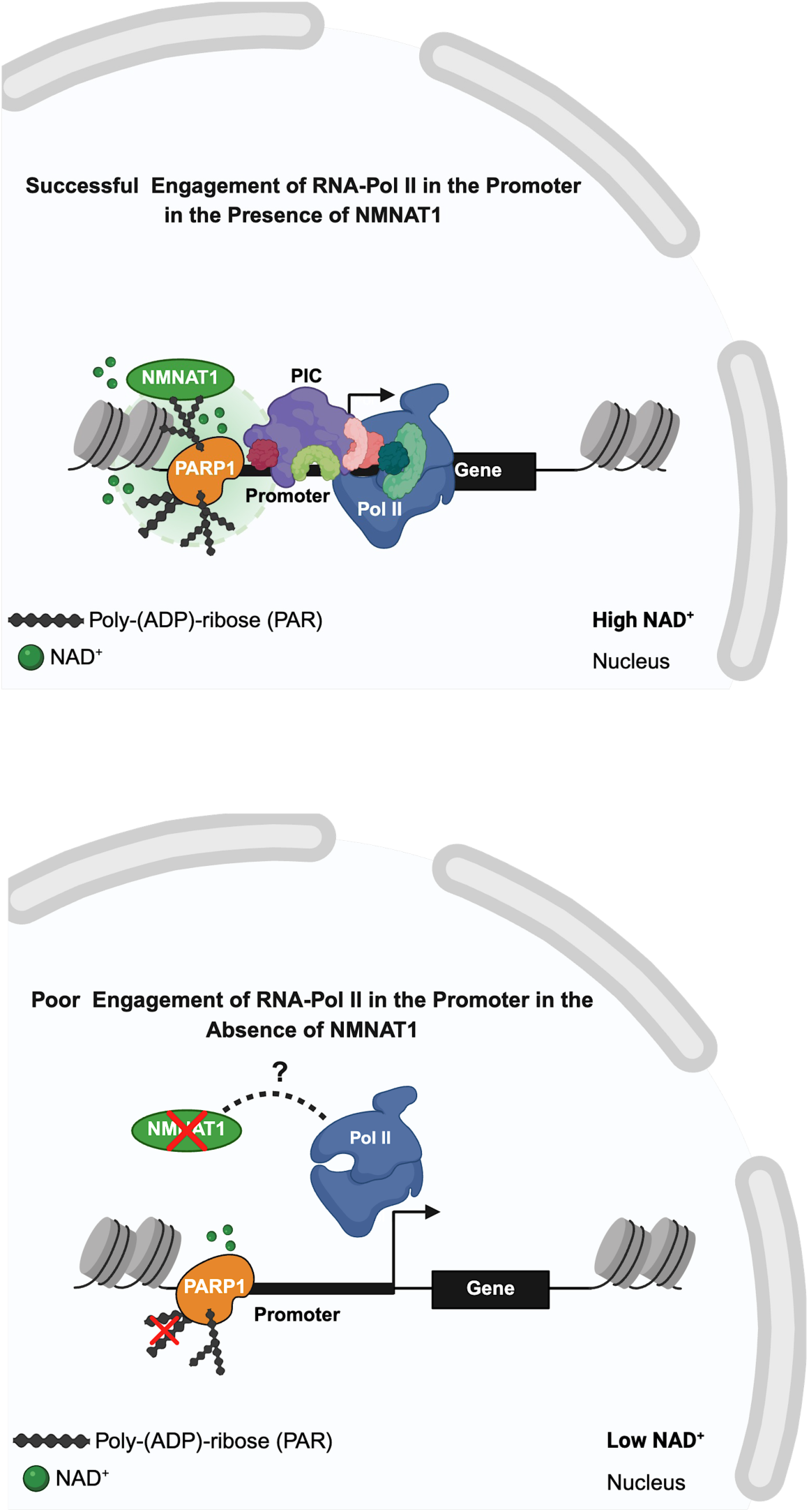
Model: Loss of NMNAT1 Impairs RNA-Pol II Recruitment through NAD⁺-Dependent Regulation of PARP1. Model: Depletion of NMNAT1 disrupts local nuclear NAD⁺ synthesis, impairing PARP1 activity and leading to poor engagement of RNA-Pol II at gene promoters. Previous models proposed that PARP1 facilitates promoter-proximal pause release of RNA-Pol II via PARylation of NELF; however, our data reveal that NMNAT1^KO^ cells exhibit reduced RNA-Pol II occupancy at transcription start sites of downregulated genes, indicating a defect in RNA-Pol II recruitment or stabilization. This suggests that NMNAT1-derived NAD⁺ may enable PARP1 to modify additional chromatin-associated factors required for RNA-Pol II engagement.

Our findings conceptually expand the understanding of transcriptional regulation by showing that a typical metabolic enzyme can directly influence transcription at distinct regulatory sites in the genome. NMNAT1 is part of a growing group of metabolite-producing enzymes that regulate gene by acting locally at promoters and enhancers. This research opens avenues for future studies on how other nuclear metabolic enzymes may similarly affect transcription. This raises new questions about how metabolism is spatially organized within the nucleus, particularly regarding the links between localized NAD⁺ production, the recruitment of transcription factors, and the formation of regulatory complexes.

Although our primary emphasis is on mechanistically dissecting transcription, these results could also provide a fresh perspective for examining NAD⁺ depletion linked to diseases. Reduced NAD⁺ levels are associated with aging, cancer, and metabolic disorders, all marked by extensive transcriptional changes. By understanding how NMNAT1 coordinates metabolic and transcriptional regulation, we can uncover how disruptions in nuclear metabolism may lead to gene dysregulation in diseases. However, clarifying these disease associations will necessitate additional research in relevant physiological models.

In summary, we define NMNAT1 as a chromatin-associated regulator of gene expression, linking NAD⁺ metabolism to RNA Polymerase II engagement at key regulatory genes. These findings highlight a critical intersection between metabolism and transcription, suggesting that local enzymatic activity is a key regulatory layer in gene expression. This work opens new conceptual directions in the transcription field, highlighting the underappreciated regulatory roles of nuclear metabolic enzymes in shaping gene expression programs.

## RESOURCE AVAILABILITY

### Lead Contact

Further information and requests for resources and reagents should be directed to and will be fulfilled by the Lead Contact, Robert A. Coleman

### Materials Availability

Materials are available upon request.

### Data and Code Availability

RNA-seq and CUT&Tag datasets have been deposited at GEO (GSE295593 and GSE295596) and are publicly available as of the publication date. Accession numbers are listed in the key resources table. The DOI is listed in the key resources table. The lead contact will share any additional information required to reanalyze the data reported in this paper upon request.

## Acknowledgments

We thank the Coleman lab members for their comments supporting this study. We would also like to thank the Einstein Epigenomics Core and Flow Cytometry Core Facilities for technical support and instrumentation. This work was supported by grants from the National Institutes of Health (NIH), including NIGMS R01GM126045 and the Chan Zuckerberg Initiative Metabolism Network (MET-0000000459) awarded to R.A.C. Y.C.C. was supported by a predoctoral fellowship from the NIH (NIGMS F31GM146429). S.J.T. was supported by a Young Investigator Award from the Edward P. Evans Foundation. M.J.G. was supported by R01 CA155232. U.S. was supported by NIH grant R35CA253127. U.S. holds the Edward P. Evans Endowed Professorship in Myelodysplastic Syndromes at Albert Einstein College of Medicine. The Endowed Professorship was supported by a grant from the Edward P. Evans Foundation. Some panels of Figures 1, 3, 4, 5, and 6 were created using BioRender.com.

## Author contributions

Conceptualization, Y.C.C. and R.A.C.; Methodology, Y.C.C., S.J.T., M.J.G. and R.A.C.; Investigation, Y.C.C. and S.J.T.; Writing – Original Draft, Y.C.C. and R.A.C.; Writing – Review & Editing, Y.C.C., R.A.C. and S.J.T.; Funding Acquisition, R.A.C. and Y.C.C.; Resources, R.A.C. U.S. and S.J.T.; Supervision, R.A.C.

## Supplementary Information

Document S1: Figure S1 and Figure S2

Table S1: Excel File containing additional RNA-seq differential expression analysis data too large to fit in a PDF, related to Figure 2.

## STAR METHODS

### EXPERIMENTAL MODEL AND SUBJECT DETAILS

#### Cell Lines

Osteosarcoma U-2OS cells (female, 15 years) were obtained from the American Type Cell Culture (ATCC, HTB-96) and were regularly tested and determined to be mycoplasma-free. Cells were cultured in high-glucose Dulbecco’s modified Eagle’s medium DMEM (1X) + GlutaMAX^TM^ (10569-010, gibco), containing 10% fetal bovine serum (S11150, R&D Systems), and 1% penicillin-streptomycin (15140-122, gibco, Life Technology) at 37°C and 5% CO_2_.

#### CRISPR/Cas9-mediated knockout of NMNAT1 in U-2OS cells

An NMNAT1 knockout cell line was generated using a commercially available NMNAT1 CRISPR/Cas9 knockout plasmid kit (sc-403085, Santa Cruz Biotechnology, Inc.) and an NMNAT1 HDR plasmid (sc-403085-HDR, Santa Cruz Biotechnology, Inc.). Cells were seeded at 60-80% confluency in a 6-well plate 24 hours prior to transfection. Cells were transfected using Lipofectamine 3000 (L3000-015, Invitrogen). The final concentration of transfected DNA was 0.5 µg at a 1:1 ratio of NMNAT1 KO plasmid to NMNAT1 HDR. Cells were incubated for 24–72 hours at 37°C and 5% CO2. Transfection efficiency was visually confirmed by the detection of red fluorescent protein (RFP) using fluorescent microscopy. To generate a stable NMNAT1 knockout cell line, we performed puromycin selection (1.3 µg/mL) for 3 weeks. After puromycin selection, cells were sorted by RFP fluorescence via FACS using a BD FACSAria III flow cytometer. The NMNAT1 mRNA and protein expressions were validated by qRT-PCR and Western blot, respectively.

### METHOD DETAILS

#### RIPA Protein Extraction and BCA Protein Quantification

Whole-cell protein extraction was performed using RIPA Lysis and Extraction Buffer (89900, Thermo Scientific). Briefly, cell pellets were washed twice in cold 1X PBS (114-056-101, Quality Biological, Inc.) and centrifuged at 500 x g for 5 minutes at 4°C. Then, the pellet was resuspended in 100 mL of RIPA Buffer containing 1X of Halt^TM^ Protease Inhibitor Cocktail (1861748, Thermo Scientific). Cells were incubated on ice for 5 minutes and then centrifuged at 14,000 x g for 15 minutes at 4°C. Protein quantification was performed using Pierce BCA Protein Assay Kit (23225, Thermo Scientific). Absorbance was measured at 564nm using SpectraMaxÒ iD3 (Molecular Devices) microplate reader and SoftMaxÒ Pro v7.1 Software (Molecular Devices).

#### Western Blotting

Protein samples were prepared with 1X Bolt LDS Sample Buffer (B0007, Novex by Life Technologies) and 2.5% β-mercaptoethanol (M3148-100ML, Sigma-Aldrich) and boiled for 5 minutes at 95-100°C. Samples were run in a 4-12% SDS-PAGE gel (NW04122BOX, Invitrogen) for 2 hours at 100V. Proteins were transferred to a nitrocellulose membrane 0.45 µm (10600003, cytiva) overnight at 4°C and 30V. Membranes were blocked for 1 hour using 5% Blotting-Grade Blocker (1706404, Bio-Rad) in 1X Tris-Buffered Saline (J60764.K2, Thermo Scientific) containing 0.1% Tween-20 (655204-100ML, Millipore) (TBS-T). Primary monoclonal antibodies including anti-NMNAT1 (1:1000, sc-271557, Santa Cruz Biotechnology, Inc.), anti-PARP1(1:1000, AB194586, abcam), anti-PAR (2.5ug/ml, ALX-804-220-R100, Enzo), anti-H3(D1H2) (1:5000, 4499, Cell Signaling Technology) were incubated overnight at 4°C in 5% milk in TBS-T. Membranes were then washed 3 times (10 minutes each) in TBS-T. Secondary monoclonal antibodies, anti-Mouse IgG Alexa Fluor 680 (1:5000, A21057, Invitrogen), anti-Rabbit IgG Alexa Fluor Plus 800 (1:5000, A32735, Invitrogen), were incubated for 1 hour at room temperature. Membranes were then washed 3 times (10 minutes each) in TBS-T. Protein bands were visualized by antibody fluorescence using a LI-COR Fc Odyssey Imaging System. Quantification was performed using Fiji image analysis software^49^.

#### Total cellular NAD^+^ measurements

Total cellular NAD^+^ concentration was measured with a colorimetric NAD^+^/NADH Cell-Based Assay Kit following the manufacturer’s instructions (No.600480, Cayman Chemical, USA). Cells were seeded in a 96-well plate at a 10^3^-10^4^ cells/well density in 120 mL culture media and incubated overnight at 37°C and 5% CO_2_. Culture media was removed, and cells were washed with Assay Buffer. Cells were then centrifuged at 500 x g for 5 minutes. The Assay Buffer was carefully aspirated to avoid disruptions of the cell monolayer. Cells were permeabilized with 110 mL Permeabilization Buffer and incubated with gentle shaking for 30 minutes at room temperature. Then, cells were centrifuged at 1,000 x g for 10 minutes at room temperature. After centrifugation, 100 mL of standards and supernatant from the cell culture wells were transferred to a new 96-well clear-bottom plate. A 100 mL of Reaction Solution was added to each well. The plate was incubated with gentle shaking for 90 minutes at room temperature. Absorbance was measured using a microplate reader, 2030 Victor X plate reader (Perkin Elmer) at 450nm. Total cellular NAD^+^ (nM) concentration was calculated using the following Equation 1.

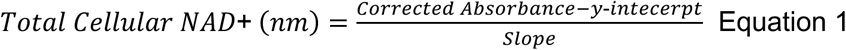

#### Total cellular ATP measurements

Total cellular ATP concentration was determined using a luminescence ATP Detection Assay Kit following the manufacturer’s instructions (No.700410, Cayman Chemical, USA). Cells were seeded in a 96-well plate at a 10^3^-10^4^ cells/well density in 200 μL culture media and incubated overnight at 37°C and 5% CO_2_. Culture media was removed, and cells were washed with pre-chilled 1X PBS (114-056-101, Quality Biological, Inc.). Then, 200 mL of ice-cold 1X ATP Detection Sample Buffer was added to the cells. Cells were homogenized by pipetting the 1X ATP Detection Sample Buffer up and down several times. Then, the cell lysate was transferred to a pre-chilled polypropylene tube. On a 96-well assay plate (3370, costar), 100 mL of Reaction Mixture and 10 mL of cell lysates were added. The plate was incubated at room temperature for 20 minutes, protected from light. Luminescence was measured with a 2030 Victor X plate reader (Perkin Elmer). Total cellular ATP was calculated using the Beer-Lambert equation with a Log transformation according to the manufacturer’s instructions (Equation 2). ATP concentration was determined by taking the antilog of Log ATP concentration (Equation 3).

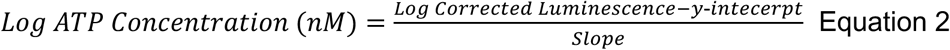

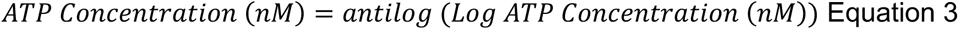

#### Calcein-AM Viability Assay

Cells were seeded into 96-well plates (3596, Corning) at a density of 2×10^4^ cells in 200mL of high-glucose Dulbecco’s modified Eagle’s medium DMEM (1X) + GlutaMAX^TM^ (10569-010, gibco), containing 10% fetal bovine serum (S11150, R&D Systems), and 1% penicillin-streptomycin (15140-122, gibco, Life Technology) and incubated overnight at 37°C and 5% CO_2_. The following day, cells were washed with 1X PBS (114-056-101, Quality Biological, Inc.) and then stained with 200 mL of Calcien-AM Fluorescent Dye (354217, Corning) to a final concentration of 1 mM in HBSS/0.01% BSA solution. Cells were incubated for 1 hour at 37°C and 5% CO_2_. Fluorescence signal was measured at Ex/Em=490/525nm using a 2030 Victor X plate reader (Perkin Elmer). Variability was expressed as a percentage of the parental control.

#### Growth Curve Assay

Cells were seeded into a 6-well plate (3596, costar) at a density of 1×10^4^ cells in 2 mL of high-glucose Dulbecco’s modified Eagle’s medium DMEM (1X) + GlutaMAX^TM^ (10569-010, gibco), containing 10% fetal bovine serum (S11150, R&D Systems), and 1% penicillin-streptomycin (15140-122, gibco, Life Technology) and incubated overnight at 37°C and 5% CO_2_. Each day, cells were trypsinized (25-052-CL, Corning), centrifuged at 1,500 rpm for 3 minutes, and then resuspended in 100 mL of culture media. Cells were stained with a 1:1 dilution of Trypan Blue Stain (0.04%) (15250-061, gibco) and counted using KOVA Plastics Glasstic Slides (87144, KOVA Plastics).

#### RNA-seq

RNA-seq was performed on U-2OS parental and NMNAT1^KO^ (polyclonal) cell lines. Cells were seeded into 150cm plates (3596, costar) at a density of 2-3×10^6^ cells in 25 mL of high-glucose Dulbecco’s modified Eagle’s medium DMEM (1X) + GlutaMAX^TM^ (10569-010, gibco), containing 10% fetal bovine serum (S11150, R&D Systems), and 1% penicillin-streptomycin (15140-122, gibco, Life Technology) and incubated overnight at 37°C and 5% CO_2_. The following day, cells were washed with 1X PBS (114-056-101, Quality Biological, Inc.) and harvested with Accutase (1000449, MP Biomedical). Then, cells were counted, centrifuged, and frozen. Samples containing 1×10^6^ cells per condition were sent to Novogene Co., LTD (Beijing, China) for RNA processing and paired-end 150-bp read length sequencing using a NovaSeq PE150 platform.

#### CUT&Tag

CUT&Tag was performed as reported in (Kaya-Okur, 2019), but with a few technical alterations. Briefly, 5 × 10^5^ cells per cell line were harvested and lightly fixed with 2% formaldehyde for 2 minutes. The cells were bound to Concanavalin A-coated beads (Bangs Laboratories) and incubated with the primary antibody (NMNAT1 (PA5-84418, Invitrogen), Pol-II Phospho-Rbp1 CTD Ser2/Ser5 (13546S, Cell Signaling Technology), or IgG control (sc-3888, Santa Cruz Biotechnology, Inc.) at 4°C overnight. Samples were then incubated with a secondary antibody (guinea pig α-rabbit (Antibodies Online, ABIN101961)), followed by adding a pre-loaded pA-Tn5 adapter complex (generated in-house). Tagmentation buffer with magnesium was used to induce transposase fragmentation. DNA was extracted by phenol/chloroform/isoamyl alcohol and amplified with NEBNext HiFi 2x PCR Master mix and universal i5 and barcoded i7 primers for 13 cycles. AMPure XP beads (A63880, Beckman Coutler) were used for post-PCR clean-up of the libraries. Libraries were subject to 35 bp paired-end sequencing using the Illumina NextSeq 500 platform with 35 bp paired-end reads on high output mode at the Einstein Epigenomics core. Fastq files were generated using Picard tools v2.17.1 with adapter trimming by TrimGalore v0.3.7 and QC assessment using FASTQC v0.11.4.

### QUANTIFICATION AND STATISTICAL ANALYSIS

#### RNAseq Analysis

RNAseq-generated fastq files were examined for quality control by FastQC. Adapters were trimmed using Trimgalore v0.3.7, and FastQC v0.11.4 was used to analyze base sequence quality, GC content, N content, and sequencing duplication levels. Reads were mapped to the human hg38 transcriptome using the STAR aligner^50^. Then, raw counts were normalized and analyzed for differential expression in R using the Bioconductor package DESeq2^51^. Differentially expressed genes were assigned an adjusted p-value <0.05 by DESeq2. For GSEA analysis, pre-ranked gene lists were generated using the negative logarithm of the adjusted p-value multiplied by the sign of the fold change for each gene (Equation 4) and input into GSEA Pre-ranked to calculate the enrichment score for each gene set^52^.

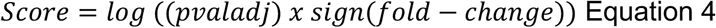

Pre-ranked gene lists were queried against standard MSigDB gene sets, including hallmark and c1-7 curated gene sets. In addition, Gene Ontology Terms analysis and plots were generated using the ClusterProfiler^53^ package in R.

#### NMNAT1 CUT&Tag Analysis

CUT&Tag generated fastq files were mapped to the human genome (hg38) using bowtie2 (Version 2.2.3) with options: --end-to-end --very-sensitive --no-mixed --no-discordant --phred33 - I 10 -X 700. The NMNAT1 background signal of the NMNAT1^KO^ cell line was subtracted from the NMNAT1 U-2OS control cell line using deepTools2^54^ bamCompare with options –operation subtract, –normalizeUsing RPKM –binSize 50, and –outFileFormat bedgraph. Peak calling was performed using bdgpeakcall from MACS2^55^ (Version 2.1.0) with options -l 200, -g 100, and -c 5.6. Bigwig files were generated using bedGraphToBigWig for visualization in IGV_2.16.0^56^. Motif analysis and peak annotation were performed using the Homer package^57^. Meta plots were created using deepTools2^54^.

#### RNA-Pol II Ser5 CUT&Tag Analysis

CUT&Tag generated fastq files were mapped to the human genome (hg38) using bowtie2 (Version 2.2.3) with options: --end-to-end --very-sensitive --no-mixed --no-discordant --phred33 - I 10 -X 700. Normalized bedgraph files were generated using bedtools^58^ genomecov with a normalization factor of 1,000,000/No. total human reads. Peak calling was performed using bdgpeakcall from MACS2^55^ (Version 2.1.0) with options -l 100 and -c 2. Bigwig files were generated using bedGraphToBigWig for visualization in IGV_2.16.0^56^. Motif analysis and peak annotation were performed using the Homer package^57^. Meta plots were created using deepTools2^54^. Differential peak analysis was performed in R Studio with a custom script termed “GoodpeaksScript” (https://github.com/steidl-lab/rePU.1sitioning) if n=1. For GoodpeakAnalysis, three stringent filters were used for the differential peak analysis of the average peak intensity: minimum intensity of >10, minimum fold change of >2, and minimum summit of >2.

#### RNA-seq and CUT&Tag Integrated Data Analysis

Differentially expressed genes (DEGs) and identified peaks (NMNAT1 or RNA-Pol II Ser5) were integrated using either bedtools^58^ intersect v2.30.0 or a custom R script using the ChIPseeker^59^ v1.42.1. To assess the statistical association between NMNAT1 or RNA-Pol II binding and differential gene expression, we performed Fisher’s Exact Test using a 2×2 contingency table comparing the presence or absence of NMNAT1/RNA-Pol II Ser5 peaks with gene expression categories (upregulated, downregulated). Log₂ odds ratios and *p*-values were calculated to evaluate enrichment or depletion. Significance was defined as *p* < 0.05. Analyses were performed in R Studio using the fisher.test function and results were visualized using custom scripts in R Studio. Gene Ontology analysis and plots were generated using the ClusterProfiler^53^ package in R.

#### Enhancer Occupancy and Functional Annotation of NMNAT1

Enhancer regions specific to the U-2OS cell line were obtained from EnhancerAtlas 2.0^60^ and mapped to the hg38 genome assembly. Only high-confidence enhancer annotations (prediction score ≥ 0.7) were retained for analysis. Enhancers were annotated with the Homer package^57^. We then integrated enhancer annotations with RNA-seq DESeq data to determine differentially expressed genes associated with enhancers. Then, we used the DEGs-associated enhancer gene list to identify NMNAT1 binding using the enhancer coordinates using bedTools^58^ intersect v2.30.0, requiring a minimum 1 bp overlap. Enhancer-bound peaks were categorized into TSS-associated enhancers (±1 kb of TSS), gene body enhancers (within exons/introns but outside the TSS window), and intergenic enhancers (>1 kb from any gene). To evaluate whether NMNAT1 binding at enhancers was associated with transcriptional changes, enhancer-associated genes were compared to DEGs identified from RNA-seq. Fisher’s Exact Test and log₂ odds ratios were calculated using the fisher.test function in R Studio to determine statistical enrichment among DEG categories.

#### Statistical Analysis

Each experiment included three biological replicates unless otherwise specified in the Methods. Statistical analyses were performed using GraphPad Prism 10. Unpaired t-tests were used to compare two groups; one-way ANOVA with appropriate post-hoc testing was applied for multiple groups. A significance threshold of *p* < 0.05 was used throughout. Fisher’s Exact Test was performed for genomic overlap analyses to assess statistical enrichment of CUT&Tag peaks among differentially expressed genes. The respective Methods sections describe additional statistical approaches specific to high-throughput datasets.

