## Supplementary material for "NMNAT1 Binding at Promoters and Enhancers Couples NAD^+^ Synthesis to RNA Polymerase II Engagement": Suplemental Figures

**Figure S1**

**A**

| **Fisher's Exact Test-NMNAT1-Bound at Promoter-TSS** | | | |
|---|---|---|---|
| | **NMNAT1 Peaks** | **No NMNAT1 Peaks** | **Total** |
| **Up DEGs** | 659 | 862 | 1,521 |
| **Unchanged** | 5,835 | 6,746 | 12,581 |
| **Total** | 6,494 | 7,608 | 14,102 |
| **Odd Ratio** | | **0.883** | |
| **Log2 Odd Ratio** | | **-0.178** | |
| **p-value** | | **2.552e-02** | |

**B**

| **Fisher's Exact Test-NMNAT1-Bound at Promoter-TSS** | | | |
|---|---|---|---|
| | **NMNAT1 Peaks** | **No NMNAT1 Peaks** | **Total** |
| **Down DEGs** | 921 | 662 | 1,583 |
| **Unchanged** | 5,573 | 6,946 | 12,519 |
| **Total** | 6,494 | 7,608 | 14,102 |
| **Odd Ratio** | | **1.733** | |
| **Log2 Odd Ratio** | | **0.794** | |
| **p-value** | | **1.309e-24** | |

**C**

| **Fisher's Exact Test-NMNAT1-Bound at Enhancers** | | | |
|---|---|---|---|
| | **NMNAT1 Peaks** | **No NMNAT1 Peaks** | **Total** |
| **Up DEGs** | 242 | 1,279 | 1,521 |
| **Unchanged** | 2,040 | 10,541 | 12,581 |
| **Total** | 2,282 | 11,820 | 14,102 |
| **Odd Ratio** | | **0.977** | |
| **Log2 Odd Ratio** | | **-0.032** | |
| **p-value** | | **0.796** | |

**D**

| **Fisher's Exact Test-NMNAT1-Bound at Enhancers** | | | |
|---|---|---|---|
| | **NMNAT1 Peaks** | **No NMNAT1 Peaks** | **Total** |
| **Down DEGs** | 356 | 1,227 | 1,583 |
| **Unchanged** | 1,926 | 10,593 | 12,519 |
| **Total** | 2,282 | 11,820 | 14,102 |
| **Odd Ratio** | | **1.595** | |
| **Log2 Odd Ratio** | | **0.674** | |
| **p-value** | | **3.786e-12** | |

**Figure S1. Statistical Analysis for NMNAT1 Chromatin Occupancy at Promoters and Enhancers of Differentially Expressed Genes.**

Fisher's exact tests quantifying the association between NMNAT1 binding and transcriptional changes after NMNAT1 knockout in U-2OS cells. (A–B) Fisher's tests for NMNAT1 peaks at promoter–TSS regions, compared to transcriptionally upregulated (A) or downregulated (B) differentially expressed genes (DEGs). (C–D) Fisher's tests for NMNAT1 peaks at enhancer regions, compared to upregulated (C) or downregulated (D) DEGs. DEGs were defined by RNA-seq with adjusted p-value < 0.05. Odds ratios, $\log_2$ odds ratios, and p-values are shown. See also Figure 4C–D.

Figure S2

**A**

| Fisher's Exact Test-RNA-Pol II Decreased Peaks | RNA-Pol II Peaks | No RNA-Pol II Peaks | Total |
|---|---|---|---|
| Up DEGs | 37 | 1,484 | 1,521 |
| Unchanged | 790 | 11,791 | 12,581 |
| Total | 827 | 13,275 | 14,102 |
| Odd Ratio | | **0.372** | |
| Log2 Odd Ratio | | **-1.426** | |
| p-value | | **3.049e-11** | |

**B**

| Fisher's Exact Test-RNA-Pol II Decreased Peaks | RNA-Pol II Peaks | No RNA-Pol II Peaks | Total |
|---|---|---|---|
| Down DEGs | 297 | 1,286 | 1,583 |
| Unchanged | 530 | 11,989 | 12,519 |
| Total | 827 | 13,275 | 14,102 |
| Odd Ratio | | **5.223** | |
| Log2 Odd Ratio | | **2.384** | |
| p-value | | **2.903e-84** | |

**C**

| Fisher's Exact Test-RNA-Pol II Increased Peaks | RNA-Pol II Peaks | No RNA-Pol II Peaks | Total |
|---|---|---|---|
| Up DEGs | 240 | 1,281 | 1,521 |
| Unchanged | 889 | 11,692 | 12,581 |
| Total | 1,129 | 12,973 | 14,102 |
| Odd Ratio | | **2.463** | |
| Log2 Odd Ratio | | **1.300** | |
| p-value | | **8.255e-27** | |

**D**

| Fisher's Exact Test-RNA-Pol II Increased Peaks | RNA-Pol II Peaks | No RNA-Pol II Peaks | Total |
|---|---|---|---|
| Down DEGs | 84 | 1,499 | 1,583 |
| Unchanged | 1,045 | 11,474 | 12,519 |
| Total | 1,129 | 12,973 | 14,102 |
| Odd Ratio | | **0.615** | |
| Log2 Odd Ratio | | **-0.700** | |
| p-value | | **1.169e-05** | |

**Figure S2: Statistical Analysis for RNA Polymerase II Occupancy Changes at Differentially Expressed Genes After NMNAT1 Knockout.**

Fisher's exact tests quantifying the association between RNA Polymerase II (Pol II) occupancy and transcriptional changes after NMNAT1 knockout in U-2OS cells. (A–B) Fisher's tests for genes with decreased RNA-Pol II peaks, compared to transcriptionally upregulated (A) and downregulated (B) differentially expressed genes (DEGs). (C–D) Fisher's tests for genes with increased RNA-Pol II peaks, compared to transcriptionally upregulated (C) and downregulated (D) DEGs. RNA-seq defined DEGs with adjusted p-value < 0.05. Odds ratios, $log_2$ odds ratios, and p-values are shown. See also Figure 5E.
